# Heterologous expression of human pro-inflammatory Caspase-1 in *Saccharomyces cerevisiae* and comparison to pro-apoptotic Caspase-8

**DOI:** 10.1101/2021.02.07.430150

**Authors:** Marta Valenti, María Molina, Víctor J Cid

## Abstract

Caspases are a family of cysteine proteases that play an essential role in inflammation, apoptosis, cell death, and development. Here we delve into the effects caused by heterologous expression of human Caspase-1 in the yeast *Saccharomyces cerevisiae* and compare them to those of Caspase-8. Overexpression of both caspases in the heterologous model led to their activation, and caused mitochondrial depolarization, ROS production, damage to different organelles, and cell death. All these effects were dependent on their protease activity, and Caspase-8 was more aggressive than Caspase-1. Growth arrest could be at least partially explained by dysfunction of the actin cytoskeleton as a consequence of the processing of the yeast Bni1 formin, which we identify here as a likely direct substrate of both caspases. Through the modulation of the *GAL1* promoter by using different galactose:glucose ratios in the culture medium, we have established a scenario in which Caspase-1 is sufficiently expressed to become activated while yeast growth is not impaired. Finally, we used the yeast model to explore the role of death-fold domains (DD) of both caspases in their activity. Peculiarly, the DDs of either caspase showed an opposite involvement in its intrinsic activity, as the deletion of the caspase activation and recruitment domain (CARD) of Caspase-1 enhanced its activity, while the deletion of the death effector domain (DED) of Caspase-8 diminished it. We propose the yeast model as a useful and manageable tool to explore Caspase-1 structure-function relationships, the impact of mutations or the activity of putative inhibitors or regulators.

## Introduction

Caspases are a family of cysteine proteases that cleave their targets after aspartic acid residues, playing an essential role in inflammation, apoptosis, cell death, and development [1]. Mammalian caspases are classified into two major groups: pro-inflammatory and pro-apoptotic caspases. They are produced as zymogens that are activated by proteolysis upon diverse stimuli. Among them, Caspase-1 and Caspase-8 are two of the most deeply characterized members. Caspase-1 exerts its function as a pro-inflammatory caspase by promoting interleukin IL-1β activation and release via pyroptosis, a form of programmed cell death (PCD) [2–4]. Caspase-8 takes part in apoptotic PCD as an initiator caspase, upstream effector caspases in the extrinsic pathway [5]. Although they intervene in different signaling hubs, they share many structural features. Both caspases are composed of a Death-fold Domain (DD: CARD –CAspase Recruitment Domain– for Caspase-1; and DEDs –Death Effector Domain– for Caspase-8), a long, and a short catalytic subunit (Fig. 1(a)) [6].

**Figure 1.**
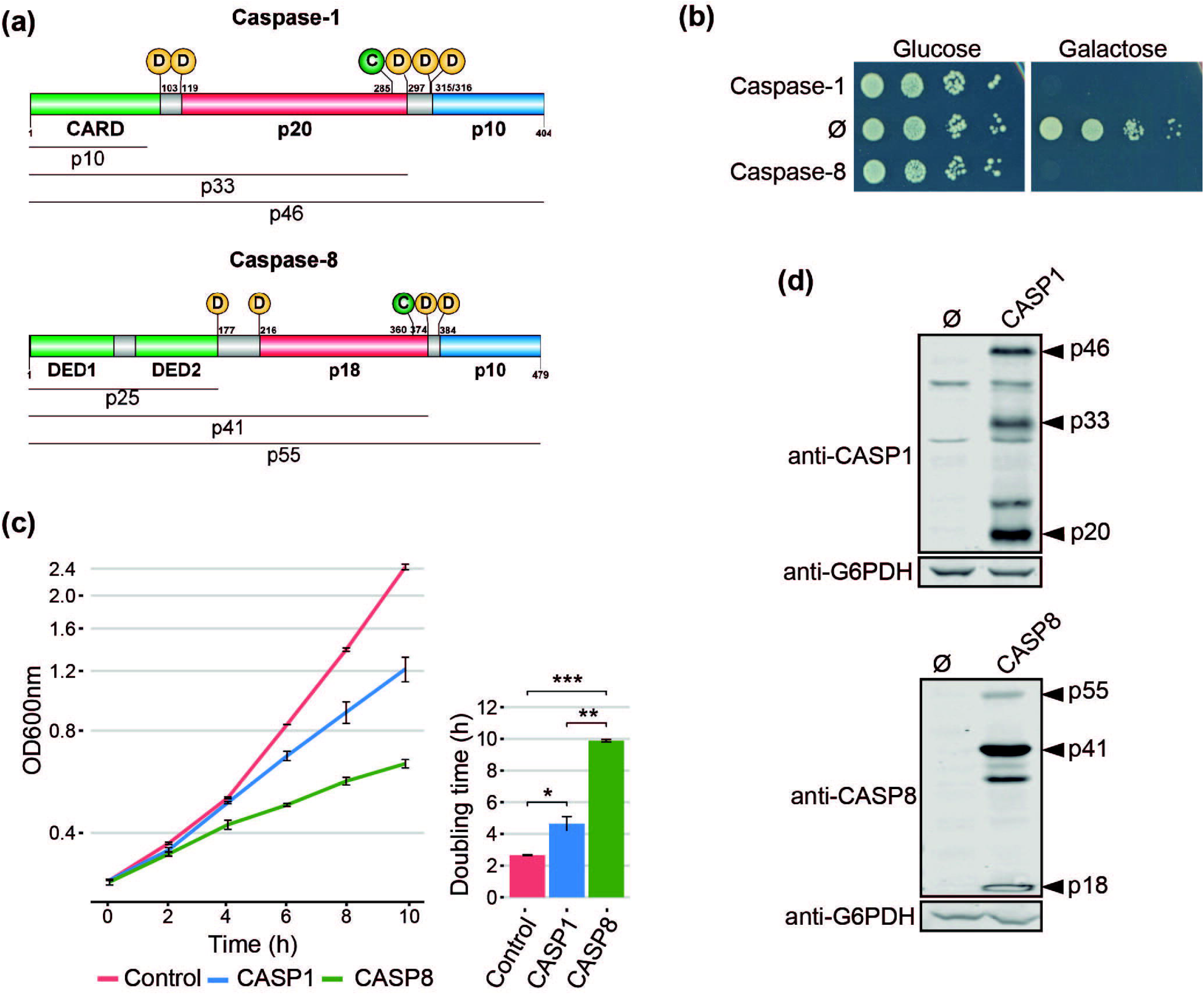
Heterologous expression of human Caspase-1 and Caspase-8 inhibits *S. cerevisiae* cell growth. (a) Schematic representation of Caspase-1 and 8 depicting their respective DDs (green), long (red), and short (blue) catalytic subunits. Their potential cleavage products and their size, the autocleavage aspartic residues (D), and the cysteine residue at the catalytic center (C) are also indicated. (b) Spot growth of BY4741 strain bearing pAG413-Caspase-1 and pAG413-Caspase-8. pAG413 empty vector (Ø) was used as a negative control. Cells were cultured on SD (Glucose) and SG (Galactose) agar media for repression and induction of Caspase-1 and Caspase-8 expression, respectively. A representative assay from three different experiments with different transformant clones is shown. (c) Growth curves of cells bearing the same plasmids as in (b) performed in SG medium. Measures of OD_600_ were taken each two hours throughout the exponential growth phase. Results are represented as OD_600_ vs time in a semilogarithmic plot (left panel). Doubling times were determined by calculating the slope over the linear portion of the growth curve (right panel). Results correspond to the mean of three biological replicates performed on different transformants. Error bars represent SD. Asterisks (*, **, ***) indicate a p-value <0.05, 0.01, and 0.01 by the Student’s T-test. (d) Immunoblots showing the expression of Caspase-1 (upper panel) and Caspase-8 (lower panel) in yeast lysates of cells bearing the same plasmids as in (b) after 5 h induction in SG medium. Membranes were hybridized with anti-Caspase-1 and anti-Caspase-8 antibodies. Anti-G6PDH antibody was used as loading control. A representative blot from three different experiments with different transformants is shown.

Under specific stimuli, Caspase-1 and Caspase-8 are recruited to macromolecular structures, known as Supramolecular-Organizing Centers (SMOCs), through heterotypic interactions between their DDs and the corresponding adaptors [7, 8]. Next, caspases dimerize and autoactivate by proteolysis. The first cleavage between the long and short catalytic subunits leads to an increase of caspase-proteolytic activity. The second cleavage, between the long subunit and the DD, releases the caspase from the SMOC and restricts its activity. Thus, active caspases transmit the signal downstream to their substrates by proteolysis. The particular SMOC to which Caspase-1 and Caspase-8 are recruited, together with target specificity, accounts for the functional divergence of these two proteins [9–11]. Caspase-1 and Caspase-8 are promiscuous enzymes and they do not recognize a strict target sequence motif. Rather, the conformational structure of the target might be more relevant for recognition than the sequence flanking the Asp residue [11–13]. Despite their importance in human disease, especially in inflammatory diseases and cancer, respective to each class of caspases, the complex nature of their regulation and activity is not yet fully elucidated. Heterologous expression in genetically tractable experimental models could help in their characterization.

For the last 40 years, the yeast *Saccharomyces cerevisiae* has proved to be useful as a model for the functional study of human proteins and signaling pathways, partly as a consequence of the development of suitable heterologous expression tools [14]. Previous reports have shown that heterologous expression of human initiator pro-apoptotic Caspase-8 and Caspase-10 is toxic to *S. cerevisiae*. On the contrary, executioner pro-apoptotic caspases are not toxic, unless co-expressed with an initiator caspase or expressed in their truncated active form [15–19], because they need to be activated via proteolysis [6]. However, there is no information available regarding the expression of pro-inflammatory caspases in yeast. These caspases deserve special attention because they are crucial in defense against pathogens, cancer, auto-immune diseases, and sepsis, as part of the innate immunity [20].

In this study, we heterologously express human Caspase-1 in yeast for the first time to our knowledge, and compare its effects with those of Caspase-8. We demonstrate that Caspase-1 autoactivates and becomes toxic in yeast when overexpressed due to its own proteolytic activity. Moreover, the reduction of the levels of expression allows us to modulate this toxicity. This model provides a novel platform to readily assess the function of human Caspase-1 mutations, its inhibitors and regulators in a manageable *in vivo* experimental setting.

## Results and Discussion

### Expression of human Caspase-1, like Caspase-8, inhibits yeast cell growth

Pro-inflammatory Caspase-1 and pro-apoptotic Caspase-8 share a similar domain architecture and distribution of proteolytic sites for autoactivation (Fig 1(a)). To study whether pro-inflammatory Caspase-1 might exert toxic effects on the yeast cell, as previously reported for initiator Caspase-8 [16, 18, 19], we cloned the cDNAs encoding for both human caspases in the same expression vector under the control of galactose-inducible *GAL1* promoter. *S. cerevisiae* cells expressing each of these caspases failed to grow on solid galactose-containing media (Fig. 1(b)), and significantly decelerated growth in liquid cultures (Fig. 1(c)). The doubling time calculated through the growth curve raised from 2.5 h for control cells to 4.5 h for cells expressing Caspase-1 and almost 10 h for cells expressing Caspase-8. Biomass after 24 h of culture in galactose-based liquid medium was reduced by 2-fold for Caspase 1- and by 6-fold by Caspase-8 as compared to the empty vector control (Fig 1(c)). Our results suggest that both human caspases become active in our model by sheer overproduction in the absence of further stimuli, probably because they self-interact bypassing the requirement for nucleating factors. This supports the proximity-driven dimerization model proposed for the pro-apoptotic initiator caspases [21], which would also extend to pro-inflammatory Caspase-1. The severe toxicity of Caspase-8 in yeast is consistent with previous reports [16, 18, 19]. The relatively milder toxicity of Caspase-1 observed in liquid culture could be due either to a lower intrinsic activity or autoactivation ability, or to a differential specificity on essential heterologous protein targets in yeast.

### Caspase-1 and Caspase-8 are self-processed in yeast

The activation of executioner pro-apoptotic caspases and pro-inflammatory caspases requires dimerization and autoproteolysis of the pro-caspase at the cleavage sites that link the long and short catalytic subunits [6]. By Western blotting analyses, we demonstrated that both caspases were efficiently expressed and self-processed in yeast into their predicted active forms, as we were able to detect protein bands corresponding to the p33 and p20 cleaved subunits for Caspase-1, and the p41 and p18 cleaved subunits for Caspase-8 (Fig. 1(d)). Thus, expression of Caspase-1 in *S. cerevisiae*, like that of Caspase-8, leads to its auto-processing and activation.

### Caspase-1 and Caspase-8 protease activity and Caspase-1 autoprocessing are essential for their toxicity in yeast

To learn whether the toxic effect caused in yeast cells by both caspases reflected their protease activity, we generated a catalytically inactive mutant for each caspase by site-directed mutagenesis, in which the cysteine residues located at their respective active centers were replaced by alanine (Caspase-1 C285A and Caspase-8 C360A). As expected, these mutant proteins neither impaired yeast growth (Fig. 2(a)), nor could we detect significant amounts of their proteolyzed subunits by Western blotting on yeast lysates (Fig. 2(b)). Thus, we conclude that caspase proteolytic activity is necessary for caspase processing, autoactivation, and toxicity in yeast. Besides, we cloned in the same vector the uncleavable Caspase-1 D5N mutant, in which the five Asp residues that allow Caspase-1 autoprocessing are mutated to Asn, an amino acid that it is not targeted by this protease [22]. As with catalytically inactive Caspase-1, yeast growth was not impaired by expression of the D5N mutant (Fig. 2(a)) and we could not detect Caspase-1 proteolyzed subunits (Fig. 2(b)), emphasizing that autoprocessing of Caspase-1 is also necessary for its activation and toxicity. However, we observed lower expression levels of the D5N mutant version as compared to wild type Caspase-1, suggesting that these mutations affect protein stability, which could also contribute to the lack of toxicity observed.

**Figure 2.**
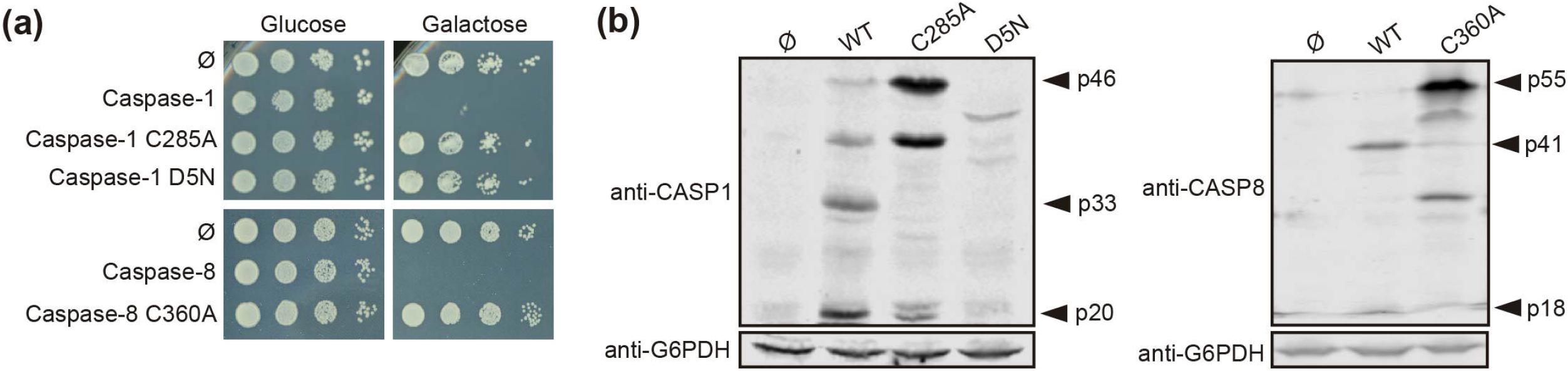
Caspase-1 and Caspase-8 toxicity is a consequence of their proteolytic activity. (a) Yeast spot assay performed as in Fig. 1(b) but using BY4741 strain bearing pAG413-Caspase-1, pAG413-Caspase-8, or plasmids bearing their respective catalytically inactive mutants pAG413-Caspase-1 C285A and pAG413-Caspase-8 C360A, and the uncleavable mutant pAG413-Caspase-1 D5N. (b) Immunoblots showing the expression in yeast lysates of cells bearing the same plasmids as in (a) after 5 h of induction in SG medium. Membranes were hybridized with anti-Caspase-1, and anti-Caspase-8 antibodies. Anti-G6PDH antibody was used as a loading control. Representative assays from three different experiments with distinct transformants are shown in all cases.

### Oxidative damage exerted by Caspase-1 in *S. cerevisiae* is less severe than that of Caspase-8

The cell death phenotype induced by pro-apoptotic initiator caspases in yeast is characterized by reactive oxygen species (ROS) production, decrease in cell viability, and propidium iodide (PI) uptake [16, 19]. To gain a better insight into the Caspase-1 terminal phenotype as compared to that of Caspase-8, we measured mitochondrial membrane depolarization and intracellular ROS production in both Caspase-1- and Caspase-8-expressing cells by staining them respectively with rhodamine 123 (Rd123) and dihydroethidium (DHE), after 5 h of induction in galactose-containing medium, and analyzing them by flow cytometry (Fig. 3(a)). We found a statistically significant increase in membrane depolarization and ROS production for both caspases as compared to control cells, which was higher with Caspase-8 than with Caspase-1. Then we evaluated whether this phenotype was accompanied by a reduction in cell viability and cell death under the same conditions. Cell viability was measured based on the ability of yeasts to form microcolonies and we observed a significant decrease in cell viability for both caspases (Fig. 3(b)). In addition, cell death was analyzed by flow cytometry using PI as an indicator of loss of membrane selective permeability. Likewise, we found a statistically significant differential increase in cell death for both caspases (Fig. 3(a)). Taken together, these results indicate that the expression of Caspase-1 and Caspase-8 leads to mitochondrial membrane depolarization and ROS production, as well as a decrease in cell viability and cell death. Furthermore, consistent with our results above, Caspase-8 effects are stronger in all cases.

**Figure 3.**
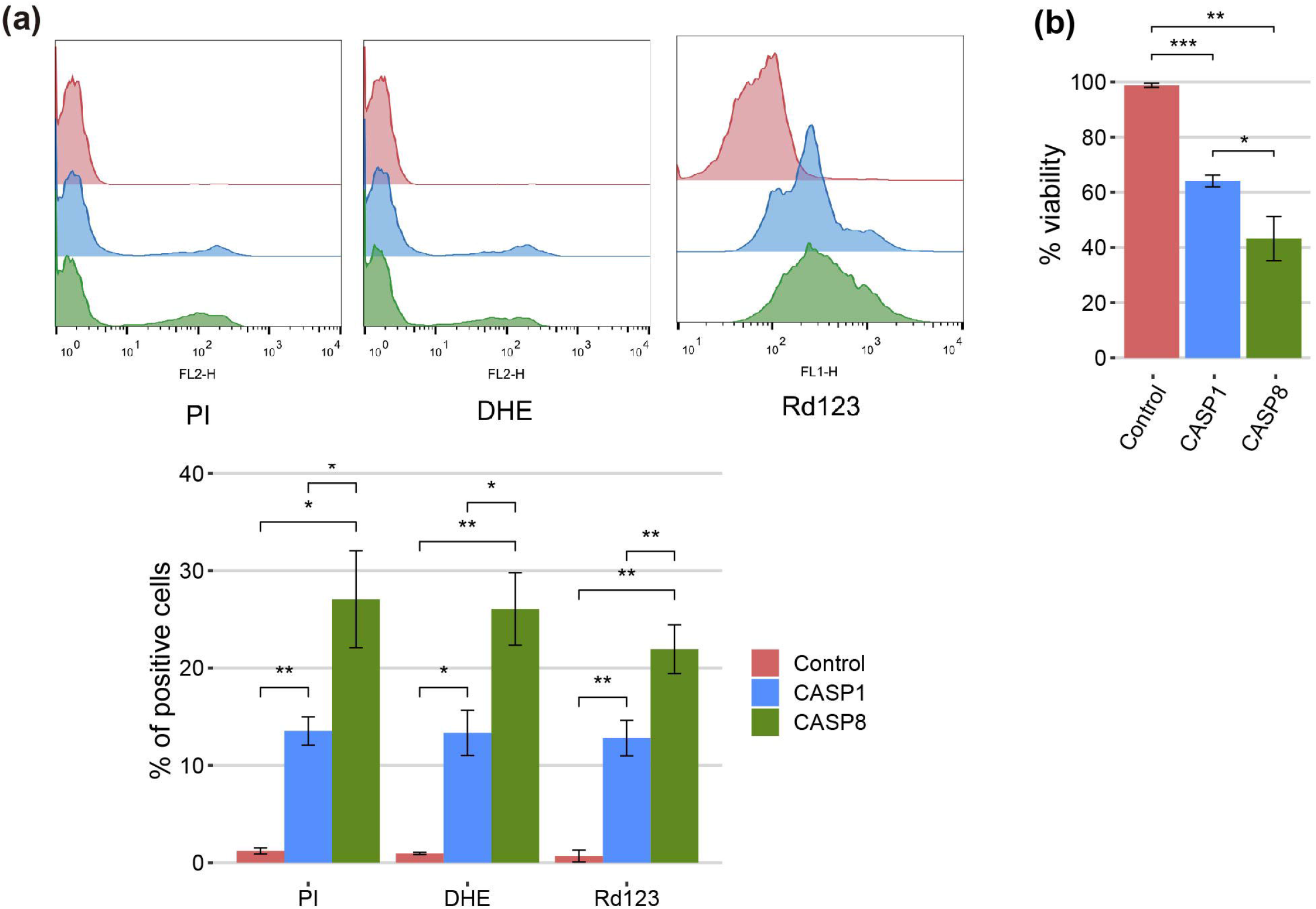
Caspase-1 and Caspase-8 cause mitochondrial membrane depolarization, ROS production, reduction in cell viability, and cell death. (a) Stacked histograms (n=10,000) showing DHE, Rd123 and PI fluorescent signal by flow cytometry respectively in abscissae (upper panel) and graph showing the percentage of positive DHE, Rd 123 and PI stained cells of each population (lower panel) of BY4741 strain bearing the same plasmids as in Fig. 1(b) after 5 h of induction in SG medium. (b) Graph showing the percentage of viable cells determined by a cell viability assay of BY4741 strain bearing the same plasmids as in (a). Results correspond to the mean of three biological replicates performed on different clones in all cases. Error bars represent SD. Asterisks (*, **,***) indicate a p-value <0.05, <0.01 and <0.001 respectively by the Student’s T-test.

### Expression of Caspase-1 and Caspase-8 differentially affects organelle morphology in yeast

We investigated the putative damages that these caspases might be causing to cellular organelles. First, to visualize mitochondrial organization, we co-expressed each caspase with an Ilv6-mCherry fusion as a mitochondrial marker. The mitochondrial tubular network was disrupted in both cases, although there were some differences between the two caspases. While mitochondria from cells expressing Caspase-1 formed large aggregates, those from cells expressing Caspase-8 were fragmented (Fig. 4(a)). Secondly, we assessed trans-Golgi network (TGN) integrity by expressing caspases in a strain marked with the TGN protein Sec7-GFP. Caspase-1 expression did not affect TGN, while Caspase-8 caused the aggregation of Golgi cisternae into one or two big spots in around 30% of the cells (Fig. 4(b)). Thirdly, we studied vacuole morphology by expressing each caspase in a strain tagged with the vacuolar protein Vph1-GFP. Both caspases caused an increment of the vacuolar diameter, although the effect was more prominent in the case of caspase-8 (Fig. 4(c)). Finally, we evaluated the endoplasmic reticulum (ER) structure by co-expressing each caspase with Sec63-mRFP as an ER marker. In this case, no differences with the control were perceived for Caspase-1-expressing cells, while 30% of Caspase-8-expressing cells displayed an expanded ER (Fig 4(d)). Previous studies report that ER expansion might be a consequence of increased ER membrane biogenesis to adapt to different stress signals [23]. In sum, expression of both caspases in yeast leads to significant organellar alterations, but the effect of caspase-8 is more severe than that of Caspase-1, especially at the ER and Golgi levels.

**Figure 4.**
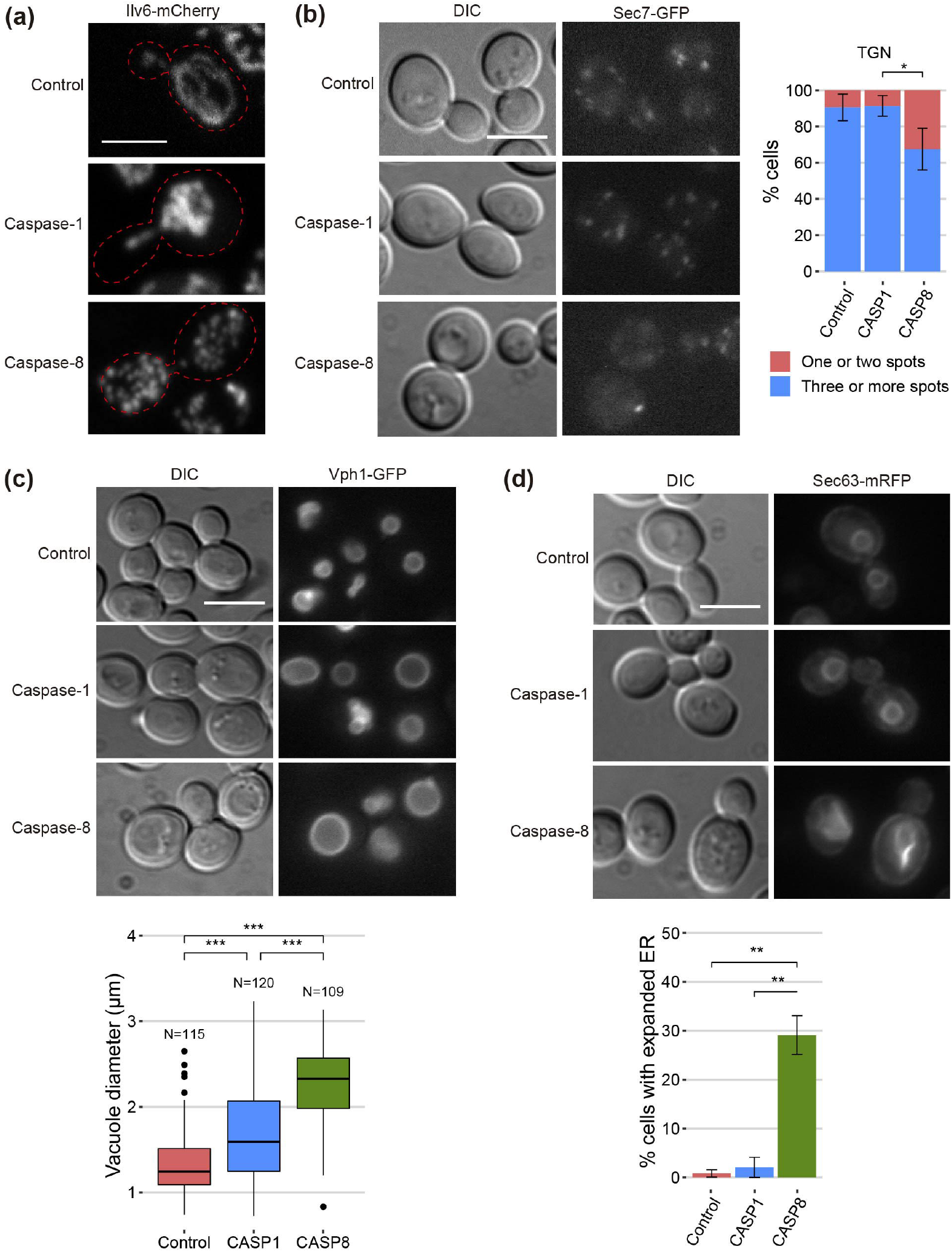
Caspase-1 and Caspase-8 overexpression alters subcellular organelles. (a) Confocal fluorescent microscopy of BY4741 *trp1Δ* strain bearing the mitochondrial marker pYEp-lac112-Ilv6-Cherry and the same plasmids as in Fig. 1(b). (b) Fluorescent and bright field differential interferential contrast (DIC) microscopy (left panel) and quantification of the number of TGN spots per cell (right panel) of DLY35 strain, bearing the TGN marker Sec7-GFP, and the same plasmids as in Fig. 1(b). (c) Fluorescent and bright field (DIC) microscopy (left panel) and boxplot of the vacuolar diameter (μm) (right panel) of MVY04 strain, bearing the vacuolar marker Vph1-GFP, and the same plasmids as in Fig. 1(b). (d) Fluorescent and bright field (DIC) microscopy (left panel) and quantification of the percentage of cells showing expanded ER (right panel) of BY4741 strain bearing the ER marker pRS425-Sec63-mRFP and the same plasmids as in Fig. 1(b). Abnormal ER expansions were considered following previously described criteria [23]. Caspase-1 and Caspase-8 expression were induced in SG medium for 5 h in all cases. All scale bars indicate 5 μm. Results in (b), (c), and (d) correspond to the mean of three biological replicates performed on different transformants. Error bars represent SD. Asterisks (*, **, ***) indicate a p-value <0.05, 0.01, and 0.001 respectively by the Student’s T-test.

### The yeast actin cytoskeleton is altered by expression of human caspases

As shown above, the dramatic growth defect observed in heterologous caspase-expressing yeast cells was accompanied by a modest loss of plasma membrane permeability, as determined by vital PI staining. This is consistent with budding arrest rather than cell lysis. However, microscopic observations did not hint a specific cell cycle arrest pattern in these cultures (data not shown), so we investigated the impact of caspases expression on the actin cytoskeleton, as it drives essential polarized secretion to promote and maintain budding. Actin staining with rhodamine-conjugated phalloidin revealed that, instead polarized patches at the growing bud, about 10% of the cells expressing any of both caspases displayed abnormal long thick actin cytoplasmic structures, which were never observed in control cultures (Fig 5(a)). These observations suggest that growth arrest in caspase-expressing yeast cells could be at least partially attributed to dysfunction of the actin cytoskeleton.

**Figure 5.**
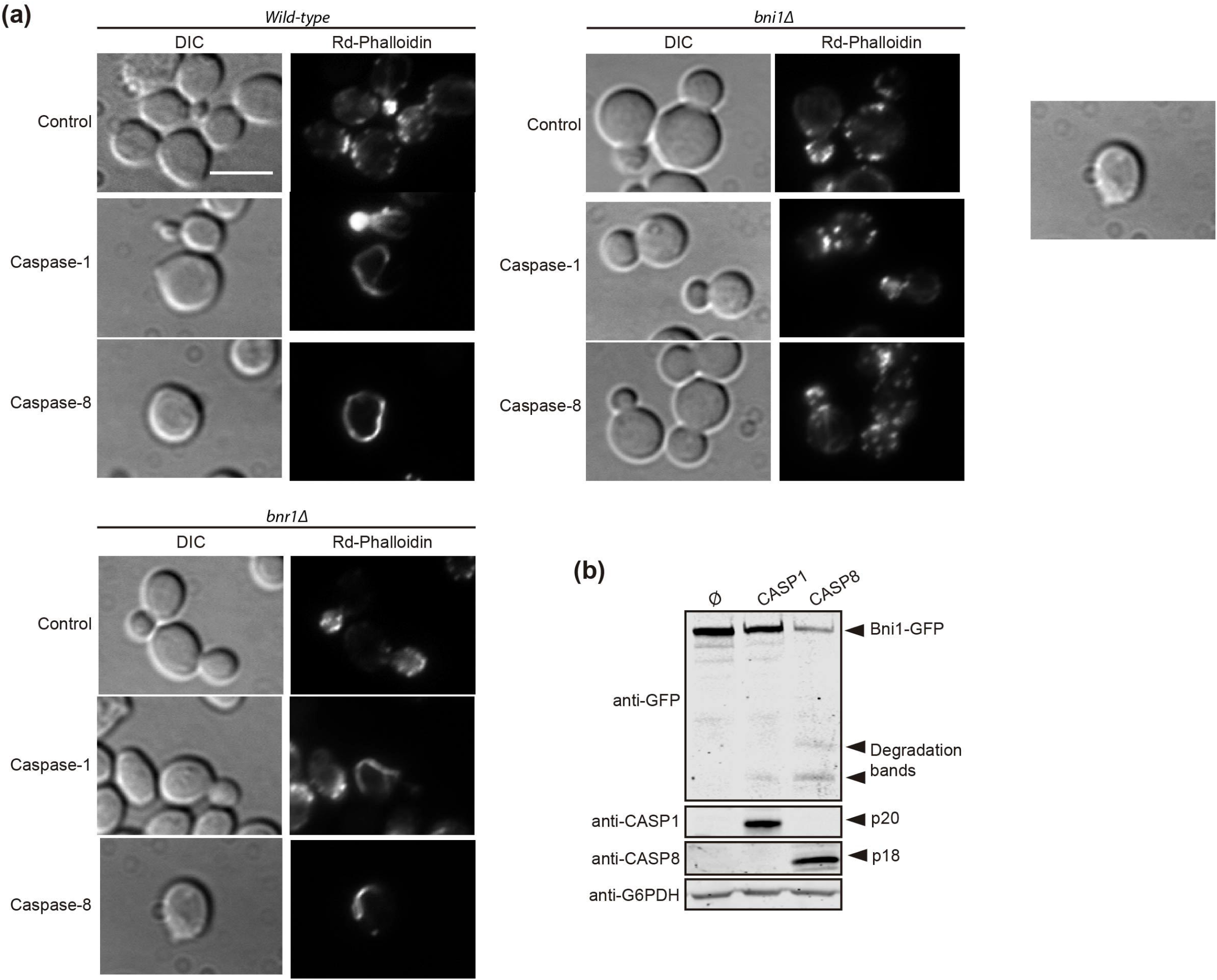
Caspase-1 and Caspase-8 overexpression alters actin cytoskeleton via Bni1 proteolysis. (a) Acting staining with rhodamine-phalloidin and bright field (DIC) microscopy of BY4741 wild type, and isogenic *bni1Δ* and *bnr1Δ* strains bearing the same plasmids as in Fig. 1(b). Scale bar indicates 5 μm. A representative image from three different experiments with distinct transformants is shown. (b) Immunoblots showing the degradation of Bni1-GFP and the expression of Caspase-1 and Caspase-8 in yeast lysates of BY4741 *Bni1-GFP-HIS2MX6* strain bearing pAG416-Caspase-1 and pAG416-Caspase-8. pAG416 empty vector (Ø) was used as a negative control. Membranes were hybridized with anti-GFP, anti-Caspase-1, and anti-Caspase-8 antibodies. Anti-G6PDH antibody was used as a loading control. Caspase-1 and Caspase-8 expression was induced in SG medium for 5 h in all cases. A representative blot from three different experiments with distinct transformants is shown.

Formins are actin-nucleating proteins that function by promoting and regulating the assembly of the actin cytoskeleton in eukaryotic cells [24]. Previous studies reported that loss of the N-terminal domain of the yeast formin Bni1 led to the formation of long cytoplasmic actin filamentous structures, resembling the ones here induced by caspases, due to uncontrolled actin polymerization caused by dysregulation of this protein [25, 26]. We hypothesized that the observed phenotype could be a consequence of Bni1 proteolysis by Caspase-1 and −8 in a region proximal to the N-terminal domain and that, in such case, the formation of these structures should be prevented in a *bni1*Δ strain. Therefore, we stained the actin cytoskeleton in caspase-expressing yeast cells individually deleted for the genes encoding each of the formins (Bni1 and Bnr1). We could observe the formation of these abnormal actin structures in the *bnr1*Δ but not on the *bni1*Δ background (Fig 5(a)), supporting our postulate that Bni1 cleavage is responsible for this phenomenon. To further confirm this hypothesis, we co-expressed a Bni1-GFP fusion with each caspase and analyzed yeast lysates by Western blotting with anti-GFP antibodies. We detected that both caspases degraded Bni1, particularly Caspase-8, as the levels of Bni1-GFP decreased and some degradation bands –absent in control lysates– appeared (Fig. 5(b)). This implies that Bni1 is a direct substrate of Caspase-1 and 8 in yeast, and that the collapse of the actin cytoskeleton into these abnormal filaments as a consequence of Bni1 cleavage likely contributes to yeast growth arrest. To our knowledge, no previous works have described any direct substrates of human caspases in yeast. Disruption of the mitochondrial network, induction of ER-phagy, or ROS production are rather unspecific damages and could account for many different causes. However, the formation of these aberrant actin structures is more specific and could be an interesting tool to assess caspase activity in yeast by microscopy or immunoblot.

### Auto-processing of Caspase-1, but not Caspase-8, can be modulated by adjusting expression levels

The development of yeast-based models for the study of human pathways should be more versatile if finely tuned regulatory mechanisms can be reproduced in the heterologous model. Thus, although a growth inhibition readout may be optimal for devising pharmacologic or genetic screens, in potential Synthetic Biology settings that imply coexpression of caspase regulators or substrates, high toxicity should be avoided. In our model, the overexpression of Caspase-1 results presumably in its dimerization and autoprocessing, thus leading to toxicity. However, we show above that its effects are not as dramatic as those caused by pro-apoptotic Caspase-8, likely due to a less efficient activation by auto-cleavage. We hypothesized that restricting Caspase-1 expression levels could prevent its dimerization and consequently reduce its toxicity in yeast. It has been reported that *GAL1* promoter induction depends on the ratio between galactose and glucose available in the culture medium, which that determines galactose uptake, rather than on the total amount of galactose. It was suggested that the competitive binding of these sugars to hexose transporters is responsible for this phenomenon [27]. Thus, we cultured Caspase-1 transformants in media containing different galactose/glucose ratios ranging from 1 to 10, always preserving a final concentration of sugars of 2%, and after 5 h of induction we analyzed Caspase-1 expression by Western blotting. As shown in Fig. 6(a), we confirmed that not only the level of expression of this protein but also its relative auto-processing capability increased gradually as the Gal/Glu ratio augmented. At those ratios for which pro-Caspase-1 was detectable but the signal for its p33 and p20 subunits was weak compared to control with galactose, caspase activity should be low. To test whether this modulation of Caspase-1 expression and autoprocessing correlated with a reduction in toxicity, we performed a spot assay using the same sugar ratios in growth media (Fig. 6(b)). Yeast growth was observed in all Gal/Glu ratios. However, in the higher ratios (8.5 and 10) we detected a reduction in the size of colonies that reflects that Caspase-1 was being processed to sufficient active form. To confirm that threshold concentrations for autoprocessing did not dramatically compromise viability, we chose R(Gal/Glu)=5.5 and R(Gal/Glu)=8.5 among the different ratios tested and analyzed yeast growth in liquid media after 24 h of induction. The 5.5 Gal/Glu ratio allowed us to express an inactive Caspase-1 (p33 and p20 bands were barely detectable in cell extracts) and the 8.5 Glu/Gal ratio led to an active Caspase-1 (p33 and p20 bands indicated the presence of the active protein). Like in solid media, although we observed a statistically significant decrease in OD_600_ due to Caspase-1 expression in both the control with galactose alone and R(Gal/Glu)=8.5, toxicity was highly reduced in the latter condition (Fig. 6(c)). Thus, this experimental setting should provide a sensitive platform for evaluating Caspase-1 activity in viable yeast cells in the future.

**Figure 6.**
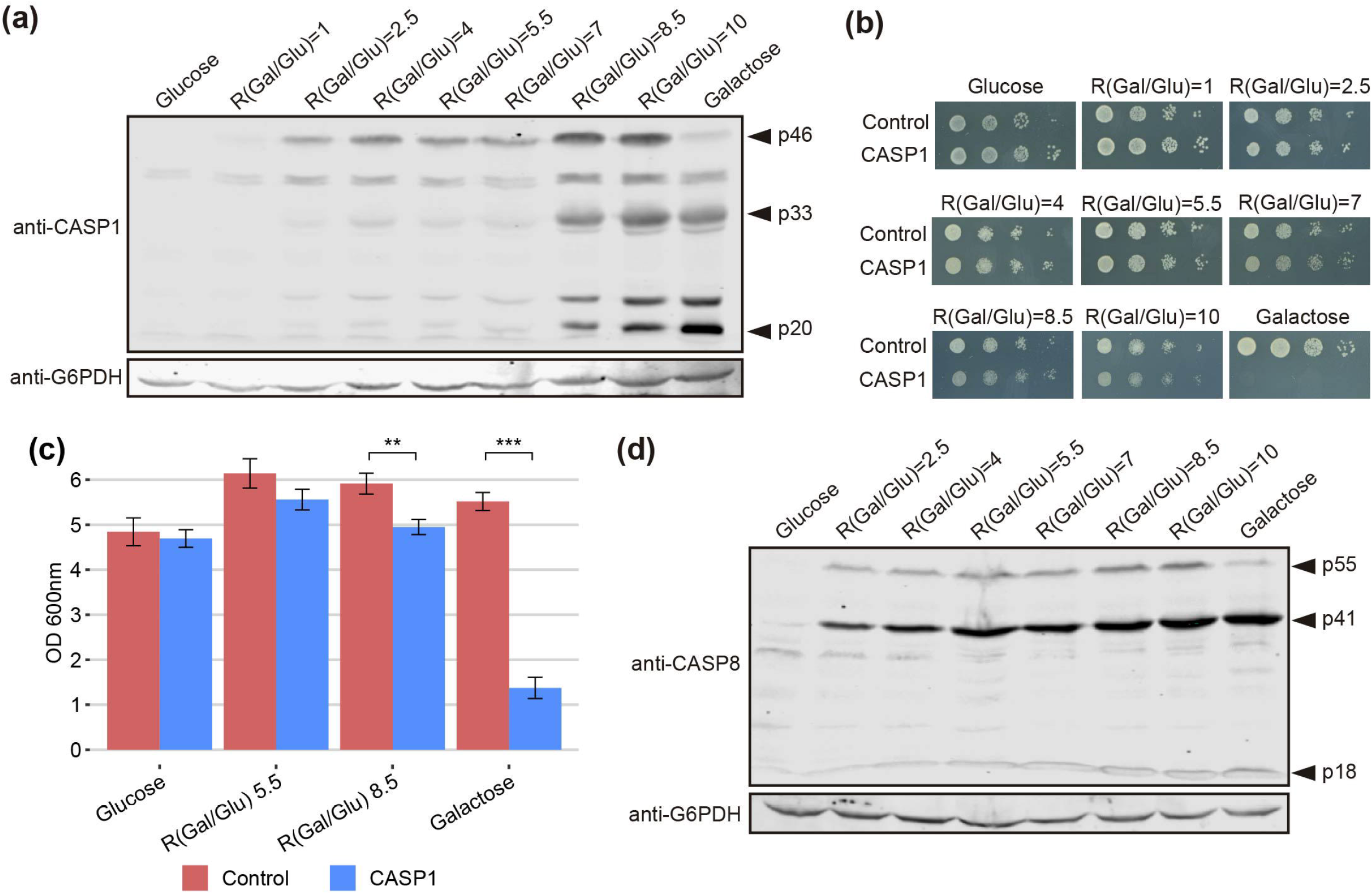
The modulation of Caspase-1 expression, but not of Caspase-8, under the *GAL1* promoter limits its auto-processing and toxicity. (a) Immunoblot showing the expression of Caspase-1 in yeast lysates of BY4741 strain bearing pAG413-Caspase-1. Cells were cultured in synthetic media containing the indicated Gal/Glu ratios with a final concentration of sugars of 2% for 5 h. Cells cultured in SD medium were used as a negative control of expression and cells cultured in SG media as a positive control. Membrane was hybridized with anti-Caspase-1 antibody. Anti-G6PDH antibody was used as a loading control. A representative assay from two different experiments with distinct transformants is shown. (b) Spot growth assay performed with the same strain and the same ratios of galactose/glucose as in (a), but in solid media. Cells bearing pAG413 empty plasmid were used as growth control. A representative assay from three different experiments with distinct transformants is shown. (c) Measurement of OD_600_ after 24 h of incubation in media containing a R(Gal/Glu)=5.5 and R(Gal/Glu)=8.5 of BY4741 strain bearing either pAG413-Caspase-1 or the pAG13 empty plasmid as growth control. As in (b), cells cultured in SD medium were used as a negative control of expression, and cells cultured in SG media as a positive control of expression. Results correspond to the mean of three biological replicates performed on different transformants. Error bars represent SD. Asterisks (**, ***) indicate a p-value <0.01, and 0.001 respectively by the Student’s T-test. (d) Immunoblotting performed as in (a) but with BY4741 strain bearing pAG413-Caspase-8. Membrane was hybridized with anti-Caspase-8 antibody. Anti-G6PDH antibody was used as a loading control. A representative assay from two different experiments with different transformants is shown.

In contrast, we could not modulate Caspase-8 activity under the same conditions tested for Caspase-1 (Fig. 6(d)) or even at lower Gal/Glu ratios ranging from 0.25 to 4 (Fig. S1). For R(Gal/Glu) ≤ 1, we could not detect Caspase-8 expression over the control in glucose, and for R(Gal/Glu) ≥ 2.5 Caspase-8 was already autoactivated.

### Deletion of Caspase-1 CARD and Caspase-8 DED domains have opposite effects

To test the strength of our model in detecting changes in caspase activity, we produced a truncated version of each caspase lacking its DD (Caspase-1 ΔCARD and Caspase-8 ΔDED, respectively). These domains facilitate the recruitment and dimerization of caspases [5, 28, 29], so we expected that their deletion would reduce protease-dependent toxicity. Contrary to these expectations, the truncated versions of these proteins were as toxic as their wild-type counterparts in a spot assay under *GAL1*-inducing conditions, as shown in Fig. 7(a), indicating that DD-mediated dimerization is not a strict prerequisite for their activation. However, when we performed PI staining after 5 h of induction in a galactose-containing medium and analyzed cells by flow cytometry, we observed opposite effects for each truncated caspase. The elimination of the CARD domain in Caspase-1 increased the percentage of PI+ cells compared to the full length caspase, while deletion of DED domain for Caspase-8 decreased the percentage of cells that lost selective permeability (Fig. 7(b)). We then checked whether these mutants were subject to autocleavage in yeast like the wild-type proteins. Indeed, we could detect by immunoblot of cell lysates the p20 subunit for Caspase-1 ΔCARD and the p18 subunit for Caspase-8 ΔDED (Fig. 7(c)).

**Figure 7.**
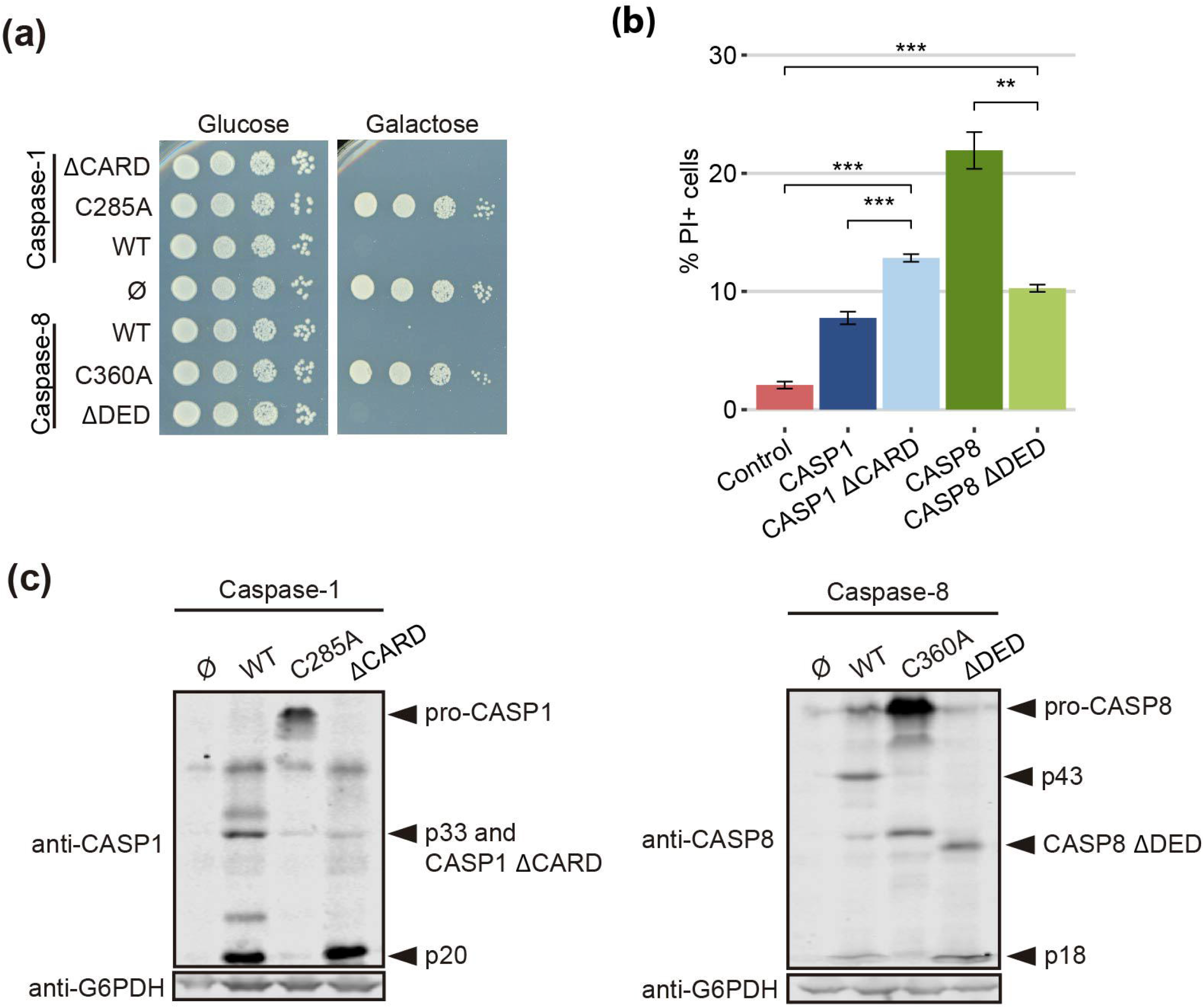
CARD and DED domains have opposite effects on the respective caspase activity. (a) Spot growth assay of BY4741 strain bearing pAG413-Caspase-1 and pAG413-Caspase-8, the plasmids with their respective catalytically inactive mutants pAG413-Caspase-1 C285A and pAG413-Caspase-8 C360A, and the plasmids with their respective mutant lacking of DD pAG413-Caspase-1 ΔCARD and pAG413-Caspase-8 ΔDED. pAG413 empty vector was used as a negative control. Cells were cultured on SD (Glucose) or SG (Galactose) agar media for induction of Caspase-1 and Caspase-8 expression. A representative assay from two different experiments with different transformants is shown. (b) Graph showing the percentage of positive PI stained cells of each population of BY4741 strain bearing the same plasmids as in (a) after 5 h of induction in SG medium. Results correspond to the mean of three biological replicates performed on different transformants. Error bars represent SD. Asterisks (**, ***) indicate a p-value <0.01, and 0.001 respectively by the Student’s T-test. (c) Immunoblots showing the expression of wild-type and the different mutants of Caspase-1 (left panel) and Caspase-8 (right panel) in yeast lysates of the cells bearing the same plasmids as in (a) after 5 h induction in SG medium. Membranes were hybridized with anti-Caspase-1 and anti-Caspase-8 antibodies. Anti-G6PDH antibody was used as a loading control. A representative assay from two different experiments with different transformants is displayed.

Next, we used the approach described in the previous section for modulating the *GAL1* promoter to verify the increment of Caspase-1 activity in the absence of its CARD domain, and the decrease of Caspase-8 activity in the absence of its DED domain. In a spot assay using ratios of galactose/glucose ranging from 1 to 10, we could observe a gradual increase of toxicity for each caspase (Fig. 8(a)). Consistent with PI staining, cell growth impairment and decrease in colony size appeared at lower ratios –corresponding to lower expression levels of the corresponding protein– for Caspase-1 ΔCARD than the full-length protein, and at higher ratios for Caspase-8 ΔDED mutant than for the corresponding fulllength caspase. Next we performed Western blotting from cells expressing Caspase-1 ΔCARD and Caspase-8 ΔDED, following an analogous strategy to the one described for Caspase-1 and Caspase-8 in Fig. 6(a,d). As shown in Fig. 8(b), Caspase-1 ΔCARD p20 cleaved subunit could be detected at very low ratios of Gal/Glu and gradually increased as the percentage of galactose raised, while this subunit could only be detected at high ratios of Gal/Glu for the full-length protein (see Fig. 6(a)). On the contrary, the cleaved p18 subunit from Caspase-8 ΔDED was detected in low levels in lysates from the increasing Gal/Glu ratios as compared to the galactose alone sample.

**Figure 8.**
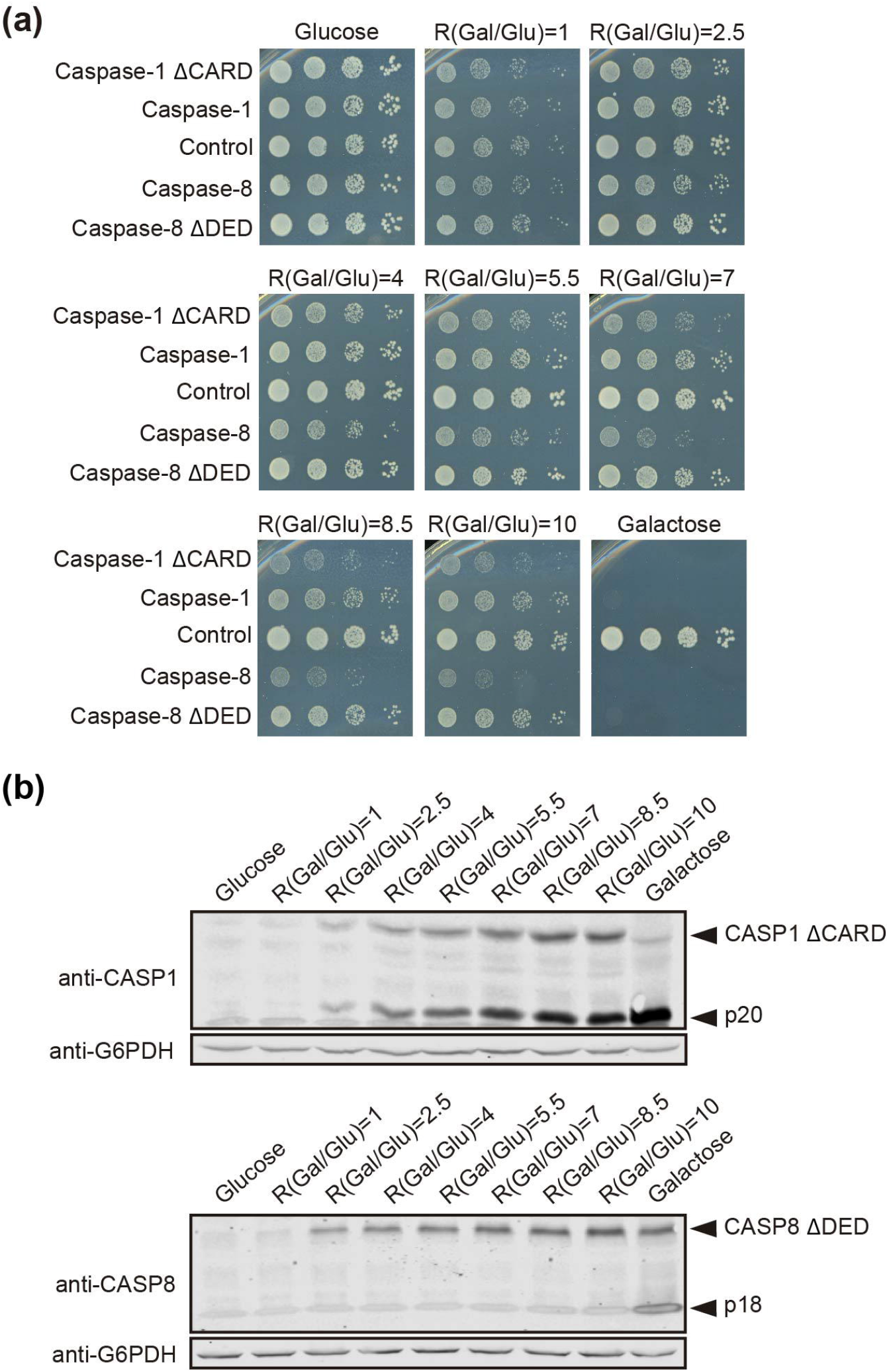
The modulation of Caspase-1 ΔCARD and Caspase-8 ΔDED expression under the *GAL1* promoter confirms their opposite effect on caspase activity. (a) Spot growth assay carried out as in Fig. 6(b) but with BY4741 strain bearing pAG413-Caspase-1 and pAG413-Caspase-8, or plasmids with their respective mutant lacking DD pAG413-Caspase-1 ΔCARD and pAG413-Caspase-8 ΔDED. pAG413 empty vector was used as a negative control. A representative assay from three different experiments with distinct transformants is shown. (b) Immunoblotting carried out as in Fig. 6(a) but with BY4741 strain bearing pAG413-Caspase-1 ΔCARD (upper panel) and pAG413-Caspase-8 ΔDED (lower panel). Membranes were hybridized with anti-Caspase-1 and anti-Caspase-8 antibody. Anti-G6PDH antibody was used as a loading control. A representative assay from two different experiments with different transformants is shown.

These results may reflect that, although DDs facilitate caspase dimerization and activation under physiological conditions by bringing closer caspase monomers to their corresponding SMOCs, at higher expression levels caspase monomers may interact with each other through other regions of the protein. Indeed, in our model the CARD domain restrains Caspase-1 activity. The proximity-driven dimerization model is compatible with the induced conformation model [21], which argues that the interaction of caspases with the SMOCs ends up in their activation because it triggers a conformational change. In this sense, CARD orientation under basal conditions may prevent Caspase-1 activation, and the interaction with its SMOC, the inflammasome, elicits a conformational change within this domain that allows Caspase-1 activation. The overexpression of a truncated version of Caspase-1 without CARD could bypass this need for a conformational change, consequently leading to a higher Caspase-1 activity. However, removal of DED reduced Caspase-8 toxicity, suggesting these DD might play different roles depending on the caspase. The physiological relevance of these results needs to be assessed in a higher eukaryote model. In human cells, caspases must be tightly regulated to preserve cell integrity, but the proteins involved in their control are probably missing in *S. cerevisiae*, as it lacks pathways closely related to those in which caspase-1 or −8 are involved. Our model provides a neat platform in which we can assess *in vivo* the intrinsic activity of caspases in the absence of other layers of regulation.

Overall, this study provides evidence that the yeast *S. cerevisiae* can be exploited as an alternative tool to study the structure, activity, and regulation of pro-inflammatory Caspase-1, as has been previously reported for Caspase-8. The expression of both caspases severely impairs yeast growth, and this readout can be appropriate for the design of pharmacologic or genetic screens. Meanwhile, the expression of sublethal concentrations of Caspase-1 with the right Galactose/Glucose ratio might be a more sensitive setting for applications such as the characterization of Caspase-1 substrates or regulators.

## Materials and methods

### Strains, media and growth conditions

The BY4741 *S. cerevisiae* strain (*MATα his3Δl leu2Δ1 met 15Δ0 ura3Δ0*) or its *trp1::kanMX4* derivative from the whole genome deletion (WGD) collection were used in all experiments unless otherwise stated. DLY35 (*MATα his3Δ1 leu2Δ0 ura3Δ0 lys2Δ0 SEC7-GFP(S65T)::KanMX*) strain (a gift from Mara C. Duncan, University of North Carolina, NC, USA)[30] was used to visualize the trans-Golgi network, and MVY04 strain (isogenic to BY4741, *VPH1-GFP-URA3*) to visualize vacuolar membrane. MVY04 strain was obtained by digesting de plasmid ZJOM153 (Addgene #268960) with *Nhe*I and *Stu*I and integrating the resulting *VPH1-GFP-URA3* fragment in the BY4741 strain. BY4741 *bnr1Δ* and BY4741 *bni1Δ* strains were obtained from the WGD collection (Euroscarf). BY4741 *Bni1-GFP-HIS3MX6* strain was obtained from the Yeast-GFP Clone Collection from UCSF. The *Escherichia coli* DH5α strain was used for routine molecular biology techniques.

Synthetic dextrose (SD) medium contained 2% glucose, 0.17% yeast nitrogen base without amino acids, 0.5% ammonium sulfate and 0.12% synthetic amino acid drop-out mixture, lacking appropriate amino acids and nucleic acid bases to maintain selection for plasmids. For synthetic galactose (SG) and synthetic raffinose (SR) media, glucose was replaced with 2% (w/v) galactose or 1.5% (w/v) raffinose, respectively. All the media components were autoclaved together. *GAL1*-driven protein induction in liquid medium was performed by growing cells in SR to mid-exponential phase and then refreshing the cultures to an OD_600_ of 0.3 directly with SG lacking the appropriate amino acids to maintain selection for plasmids for 5 h. Yeast strains were incubated at 30 °C.

For *GAL1* promoter modulation experiments, *GAL1*-driven protein induction in liquid medium and growth assays were performed as described above but instead of SG media, synthetic media containing different proportions of galactose and glucose was used. The final concentration of sugars was always 2% (w/v). In this case, the sugars were prepared at a 10x concentration and autoclaved separately from the other medium components.

### Plasmids

Transformation of *E. coli* and *S. cerevisiae* and other basic molecular biology methods were carried out using standard procedures. *CASP1* and *CASP1 ΔCARD* genes were amplified by standard PCR from pCl-Caspase-1 (a gift of Jonathan Kagan, Boston Children’s Hospital, MA, USA) using primers CASP1_Fw, CASP1 (CARD)_Fw and CASP1_Rv respectively, all designed with *attB* flanking sites. *CASP1 D5N* uncleavable mutant was amplified by standard PCR from pLEX 307-FLAG-CASP1 D5N (a gift from Daniel A. Bachovchin, Memorial Sloan Kettering Cancer Center, NY, USA) [22] using the same primers as for *CASP1. CASP8* and *CASP8 ΔDED* genes were amplified by standard PCR from pcDNA3-CASP8 (a gift from Faustino Mollinedo, CIB-CSIC, Madrid, Spain) using primers CASP8_Fw, CASP8(DED)_Fw and CASP8_Rv respectively, all designed with *attB* flanking sites. See Table 1 for primer sequences. The *attB*-flanked PCR products were cloned into pDONR221 vector by BP Gateway reaction (Invitrogen™) to generate entry clones. Subsequently, the inserts from the entry clones were subcloned into pAG413GAL-ccdB and pAG416GAL-ccdB vectors (Addgene kit #1000000011) [31] by LR Gateway reaction (Invitrogen™), generating the plasmids pGA413-Caspase-1, pAG413-Caspase-8, pAG416-Caspase-1, and pAG416-Caspase-8.

**Table 1.**
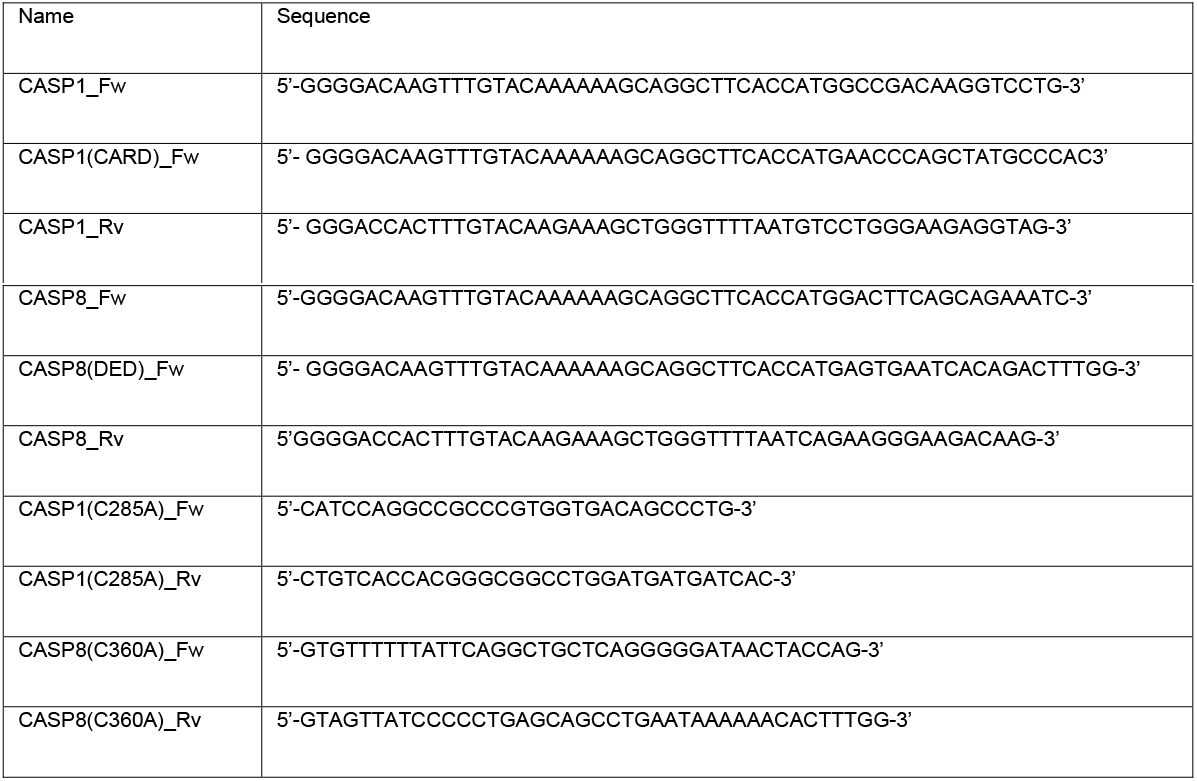
Oligonucleotides used in this work

Caspase-1 C285A and Caspase-8 C360A catalytically inactive mutants were obtained by site-directed mutagenesis performed on their respective entry clone, using primers CASP1 (C285A)_Fw and CASP1 (C285A)_Rv primers for Caspase-1 C285A and CASP8(C360A)_Fw CASP8(C360A)_Rv primers for Caspase-8 C360A. Primers are listed in Table 1. Subsequently, the inserts from the entry clones were subcloned into pAG413GAL-ccdB plasmid by LR Gateway reaction, generating the plasmids pAG413-Caspase-1 C285A and pAG413-Caspase-8 C360A.

The mitochondrial marker Ilv6-mCherry, encoded in the plasmid YEplac112-Ilv6-mCherry, has previously been described [32]. The ER marker Sec63-mRFP, encoded in plasmid pSM1959 (pRS425-Sec63-mRFP), was obtained from Addgene (#41837).

### Western blotting assays

Western blotting assays were carried out by standard techniques. Cells were harvested by centrifugation and disrupted by bead beating with a FastPrep 24 (MP Biomedicals) in 50 mM Tris-HCl pH 7.5 containing 10% glycerol, 0.1% NP-40, 1% Triton X-100, 0.1% sodium dodecyl sulfate (SDS), 150 mM NaCl, 5 mM EDTA, 50 mM NaF, 50 mM glycerol phosphate, 5 mM Na_2_P_2_O_7_, 1 mM sodium orthovanadate, 3 mM Phenylmethylsulfonyl fluoride (PMSF), and Pierce Protease Inhibitor (ThermoFisher). Lysates were cleared by centrifugation at 4°C and protein concentrations were determined by measuring the OD_280_. Proteins were resolved by sodium dodecyl sulfate polyacrylamide gel electrophoresis (SDS-PAGE) in 10% acrylamide gels, and transferred onto nitrocellulose membranes 1h at 110V. For experiments with Bni1-GFP, cells were harvested by centrifugation and disrupted with 1.85 M NaOH 7,4% β-mercaptoethanol for 10 min and trichloroacetic acid (TCA) 50% for 10 min. Cells were washed with acetone twice and resuspended in 2% SDS sample buffer. Proteins were resolved by SDS-PAGE in a 7.5% acrylamide gels and transferred onto nitrocellulose membranes overnight at 30V. Rabbit anti-Caspase-1 (D7F10) antibody (Cell Signaling Technology; 1:1000 dilution), mouse anti-Caspase-8 (1C12) (Cell Signaling Technology; 1:1000 dilution), and mouse anti-GFP (JL8) (Living colors, 1:1000 dilution) were used as primary antibodies to detect the expression of Caspase-1, Caspase-8 and proteins fused to GFP respectively. Rabbit anti-G6PDH antibody (Sigma; 1:50000 dilution) was used as a loading control. Anti-rabbit IgG-IRDye 800CW, anti-rabbit IgG-IRDye 680LT, anti-mouse IgG-IRDye 800CW, anti-mouse IG-IRDye 680LT (all from LI-COR; at 1:5000 dilution) were used as secondary antibodies. Oddissey infrared imaging system (LI-COR) was used for developing the immunoblots. Densitometry plots of were obtained using ImageJ and R.

### Flow cytometry

Cells were cultured as previously stated. After 5 h of galactose induction, 1 mL of cell culture was harvested and incubated at 30 °C with 5 μg/mL Rd 123 for 30 min in aerobic conditions, 2.5 μg/mL DHE for 5 min or 0.0005% PI for 2 min. Cells were analyzed using a FACScan (Becton Dickinson) flow cytometer through a 488 nm excitation laser and a 525/30 BP emission filter (FL1) for Rd 123 and a 585/42 BP emission filter (FL2) for DHE and PI. At least 10,000 cells were analyzed for each experiment. Data were processed using FlowJo software (FlowJo LLC, Ashland, OR, USA).

### Spot growth assays

Spot growth assays on plates were performed by incubating transformants overnight in SR media, adjusting the culture to an OD_600_ of 0.5 and spotting samples in four serial 10-fold dilutions onto the surface of SD or SG plates lacking the appropriate amino acids to maintain selection for plasmids, followed by incubation at 30 °C for 2-3 days.

### Cell viability assay

Cells were cultured as previously stated. After 5 h of galactose induction, cell viability was measured by the microcolonies method [33]. Cells suspensions (5 μL) at an adjusted OD_600_ of 0.2 were poured on a thin layer of yeast peptone dextrose (YPD) agar on a microscope slide. A coverslip was placed over the samples and after 12-24 h viable and unviable cells were identified based on their ability to form microcolonies.

### Microscopy techniques

For *in vivo* bright differential interference contrast (DIC) microscopy or fluorescence microscopy, cells were cultured as previously stated, harvested by centrifugation 3000 rpm 3 min and viewed directly on the microscope. Cells were examined with an Eclipse TE2000U microscope (Nikon) using the appropriate sets of filters. Digital images were acquired with Orca C4742-95-12ER charge-coupled device camera (Hamamatsu) and were processed with the HCImage software (Hamamatsu, Japan).

For confocal microscopy, cells were cultured as previously stated, harvested by centrifugation, and fixed with a 4% p-formaldehyde 3.4% sucrose solution for 15 min at room temperature. Then cells were washed and resuspended in phosphate buffered saline (PBS). Coverslips were washed with ethanol, treated with poly-L-lysine 0.1% solution (Sigma) for 1 h, washed with milliQ water, and dried at room temperature. Adhesion of cells was performed by adding 200 μL of fixed cells over poly-L-lysine treated coverslips and incubating for 1 h. Excess cells were removed by washing two times with PBS. ProLong™ Glass Antifade Mountant (ThermoFisher)/Glycerol (1:1) was used to avoid photobleaching. Cells were examined with an Olympus Ix83 inverted microscope, coupled to Olympus FV1200 confocal system, using the appropriate set of filters.

Observation of actin in yeast cells with rhodamine-conjugated phalloidin (Sigma) was performed as previously described [34]. For FM4-64 vital staining (*N*-[3-triethylammoniumpropyl]-4-[*p*-diethylaminophenylhexatrienyl] pyridinium dibromide; Invitrogene), cells were cultured as previously stated, harvested by centrifugation and resuspended in synthetic medium. Cells were labelled with 2.40μM FM4-64, incubated for 1.5 h at 30 °C with shaking, washed in PBS and observed by fluorescence microscopy. Images were analyzed using Image J and Adobe Photoshop.

### Statistical analysis

Statistical significance for experiments was tested with Student’s T-test. P-values were calculated with R and the asterisks (*, **, ***) in the figures correspond to a p-value of <0.05, <0.01 and <0.001 respectively. Experiments were performed as biological triplicates on different clones and data with error bars are represented as mean ± standard deviation.

## Abbreviations

DD: death-fold domain;
CARD: caspase recruitment and activation domain;
CASP1, DED: death effector domain;
PCD: programmed cell death;
SMOC: supra-molecular organizing center; Caspase-1;
CASP8: Caspase-8;
ROS: reactive oxygen species;
PI: propidium iodide
Rd123: rhodamine 123;
DHE: dihydroethidium;
TGN: trans-Golgi network;
ER: endoplasmic reticulum;
WGD: whole genome deletion;
SD: synthetic dextrose;
SG: synthetic galactose;
SR: synthetic raffinose;
OD: optical density;
SDS: sodium dodecyl sulfate;
PMSF: phenylmethylsulfonyl fluoride;
SDS-PAGE: sodium dodecyl sulfate polyacrylamide gel electrophoresis;
TCA: trichloroacetic acid;
DIC: differential interference contrast;
YPD: yeast peptone dextrose;
PBS: phosphate buffered saline

## Acknowledgements

We thank A. Sellers-Moya, D. A. Bachovchin, F. Mollinedo and M. C. Duncan for materials; and J. Kagan for materials and useful discussion; C. Mazzoni, and our colleagues at research Unit 3 for their support and discussion; and L. Sastre for technical support. M. V. was supported by a predoctoral contract from Universidad Complutense de Madrid. We thank the Genomics Unit (Genomics and Proteomics Center, UCM) for their help with the sequencing reactions, the Confocal and Multiphoton Microscopy Unit (Cytometry and Fluorescence Microscopy Center, UCM) for their help with the confocal microscopy experiments, and the Flow Cytometry Unit (Cytometry and Fluorescence Microscopy Center, UCM) for their help with the flow cytometry experiments. This research is possible thanks to funding from Grant PID2019-105342GB-I00 from Ministerio de Ciencia e Innovación (Spain).

## Notes

### Competing Interest Statement

The authors have declared no competing interest.

