## Supplemental Figure 1 for "Heterologous expression of human pro-inflammatory Caspase-1 in *Saccharomyces cerevisiae* and comparison to pro-apoptotic Caspase-8"

**Supplementary Figure S1**

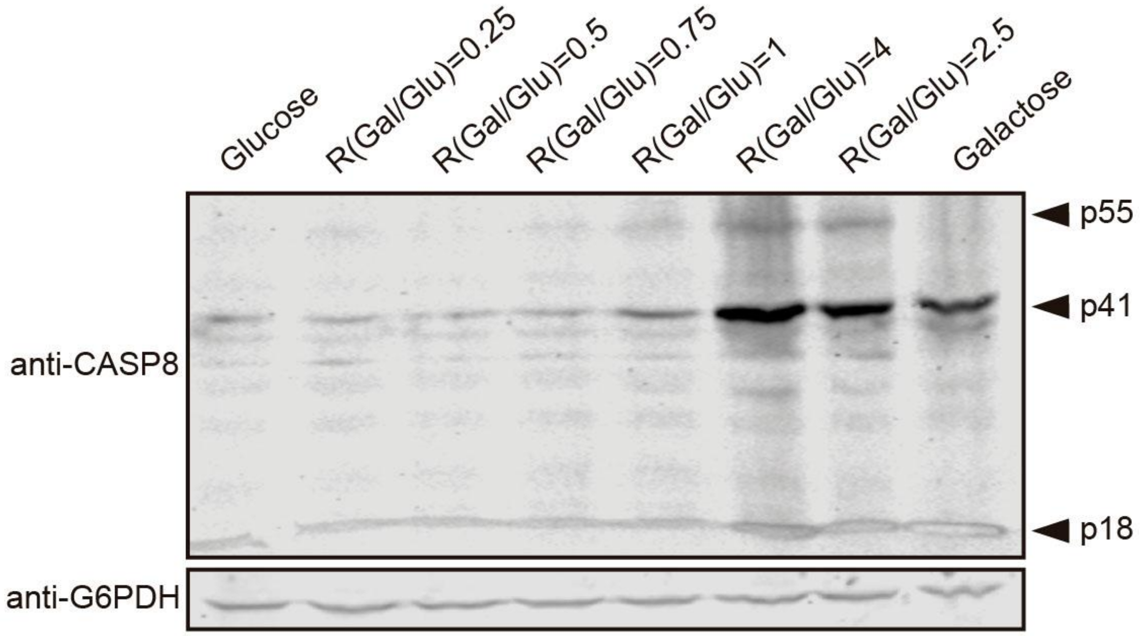


**Figure S1.** Immunoblot showing the expression of Caspase-8 in yeast lysates of BY4741 strain bearing pAG413-Caspase-8. Cells were cultured in synthetic media containing the indicated Gal/Glu ratios with a final concentration of sugars of 2% for 5 h. Cells cultured in SD medium were used as a negative control of expression and cells cultured in SG media as a positive control. Membrane was hybridized with anti-Caspase-8 antibody. Anti-G6PDH antibody was used as a loading control.
